# ATLAS: a Snakemake workflow for assembly, annotation, and genomic binning of metagenome sequence data

**DOI:** 10.1101/737528

**Authors:** Silas Kieser, Joseph Brown, Evgeny M. Zdobnov, Mirko Trajkovski, Lee Ann McCue

## Abstract

**Background:** Metagenomics and metatranscriptomics studies provide valuable insight into the composition and function of microbial populations from diverse environments, however the data processing pipelines that rely on mapping reads to gene catalogs or genome databases for cultured strains yield results that underrepresent the genes and functional potential of uncultured microbes. Recent improvements in sequence assembly methods have eased the reliance on genome databases, thereby allowing the recovery of genomes from uncultured microbes. However, configuring these tools, linking them with advanced binning and annotation tools, and maintaining provenance of the processing continues to be challenging for researchers.

**Results:** Here we present ATLAS, a software package for customizable data processing from raw sequence reads to functional and taxonomic annotations using state-of-the-art tools to assemble, annotate, quantify, and bin metagenome and metatranscriptome data. Genome-centric resolution and abundance estimates are provided for each sample in a dataset. ATLAS is written in Python and the workflow implemented in Snakemake; it operates in a Linux environment, and is compatible with Python 3.5+ and Anaconda 3+ versions. The source code for ATLAS is freely available, distributed under a BSD-3 license.

**Conclusions:** ATLAS provides a user-friendly, modular and customizable Snakemake workflow for metagenome and metatranscriptome data processing; it is easily installable with conda and maintained as open-source on GitHub at https://github.com/metagenome-atlas/atlas.

## Background

Metagenomics has transformed microbial ecology studies with the ability to generate genome sequence information from environmental samples, yielding valuable insight into the composition and functional potential of natural microbial populations from diverse environments (1, 2). Despite the prevalence of metagenome data, there are few broadly accepted standard methods, either for the generation of that data (3-5) or for its processing (6, 7). In particular, processing metagenome data in an efficient and reproducible manner is challenging because it requires implementation of several distinct tools, each designed for a specific task.

The most direct and frequently used way to analyze metagenome data is to map the sequence reads to reference genomes, when a suitable genome database from cultivated microbes is available (e.g. Humann2 (8)). However, these methods do not capture uncultivated species; studies using single-copy phylogenetic marker genes have improved estimates of species richness in metagenome data by expanding the representation of uncultivated species (9). To truly characterize a natural microbial community and examine its functional potential, assembly-based metagenome analyses are needed. This has been demonstrated by recent studies that have recovered thousands of new genomes using co-abundance patterns among samples to bin contigs into clusters (10-13).

A number of assembly-based metagenome pipelines have been developed, each providing a subset of the required tools needed to carry out a complete analysis process from raw data to annotated genomes (14-17). For example, MOCAT (16) relies on gene catalogs to evaluate the functional potential of the metagenome as a whole, but without directly relating functions to individual microbes. Anvi’o (15) requires co-assembly of the samples, which is shown to produce more fragmented assemblies (18), than assembly of individual samples. Conversely, IMP (17) permits the co-assembly of metagenomes and metatranscriptomes for individual samples, but does not allow the combination of the results. Furthermore, the configuration and technical constraints to user control often limit the adoption of these tools in the research community.

Here we present ATLAS, an assembly-based pipeline for the recovery of genes and genomes from metagenomes, that produces annotated and quantified genomes from multiple samples in one run with as little as three commands. The pipeline integrates state-of-the art tools for quality control, assembly and binning. The installation of ATLAS is automated: it depends only on the availability of Anaconda and installs all dependencies and databases on the fly. The internal use of Snakemake (19) allows efficient and automated deployment on a computing cluster.

## Implementation

The ATLAS framework organizes sequence data processing tools into four distinct analysis modules: (1) quality control, (2) assembly, (3) genome binning and (4) annotation (Fig. 1); each module can be run independently, or all four run as a complete analysis workflow. ATLAS is implemented in Python and uses the Snakemake (19) workflow manager for extensive control of external tools, including versioning of configurations and environments, provenance capabilities, and scalability on high-performance computing clusters. ATLAS uses Anaconda (20) to simplify initial deployment and environment set-up, and dependencies are handled by Bioconda (21) at runtime. Complete usage and user options are outlined in the ATLAS documentation (https://metagenome-atlas.rtfd.io).

**Figure 1.**
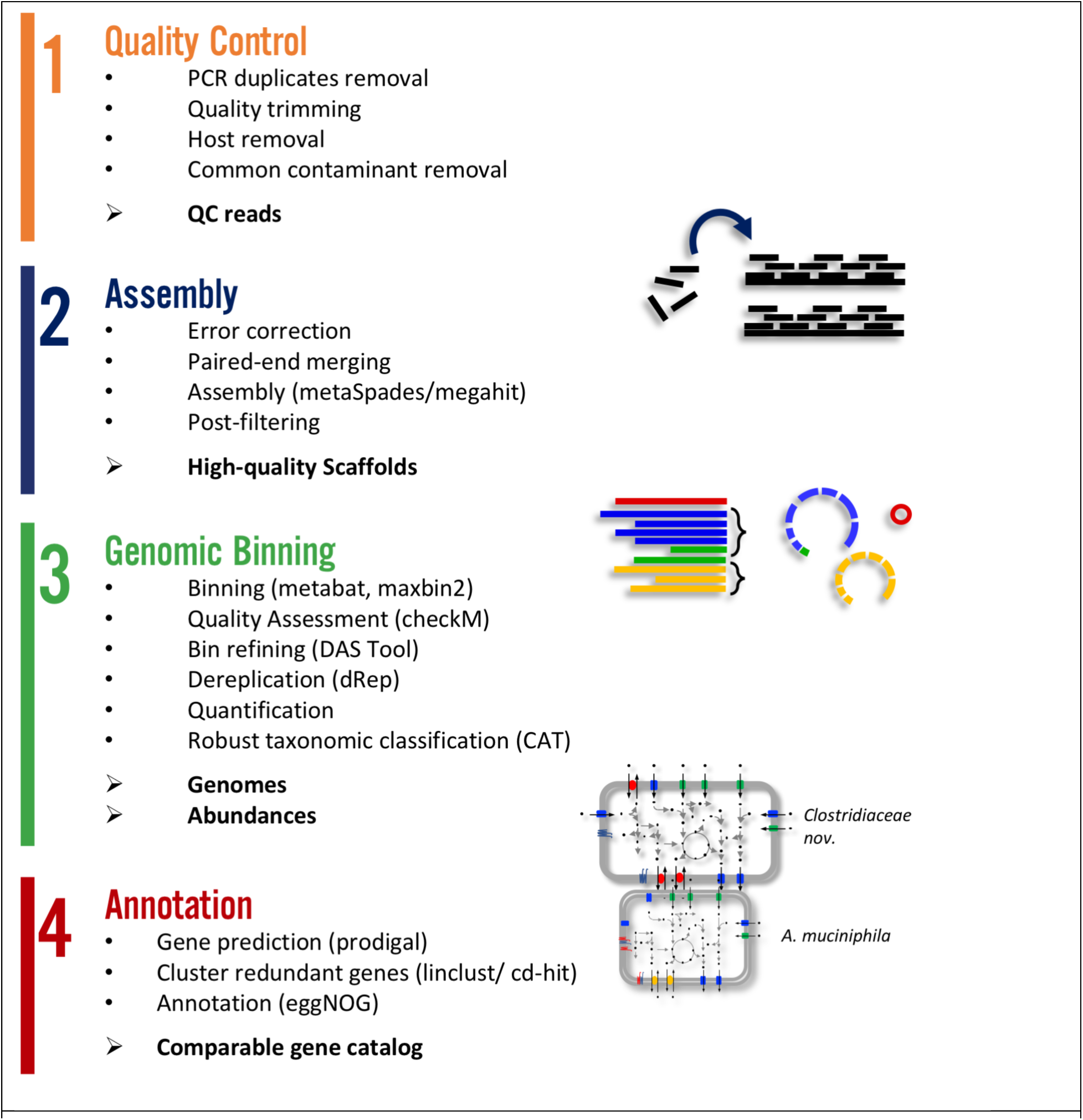
The ATLAS workflow. This high-level overview of the protocol captures the primary goal of the sub-commands that can be executed by the workflow. Individual modules can be accessed via the command line or the entire protocol can be run starting from raw sequence data in the form of single- or paired-end FASTQ files.

### Quality control

Quality control of raw sequence data, in the form of single- or paired-end FASTQ files, is performed using utilities in the BBTools suite (22). Specifically, *clumpify* is used remove PCR duplicates and (un)compress the raw data files, followed by *BBduk* to remove known adapters, trim and filter reads based on their quality and length (respectively), and error-correct overlapping paired-end reads where applicable. *BBSplit* is used to remove contaminating reads using reference sequences: PhiX is provided as a default or can be replaced by user-specified fasta-format sequences. To optimize data use, reads that lose their mate during these steps are seamlessly integrated into the later steps of the pipeline.

### Assembly

Prior to metagenome assembly, ATLAS uses additional BBTools utilities (22) to perform an efficient error correction based on k-mer coverage (*Tadpole*) and paired-end read merging (*bbmerge*). If paired-end reads do not overlap, *bbmerge* can extend them using read-derived overlapping k-mers. ATLAS supports metaSPAdes (23) or MEGAHIT (24, 25) for *de novo* assembly, with the ability to control parameters such as kmer lengths and kmer step size for each assembler. The quality-controlled reads are mapped to the assembled contigs, and bam files are generated to facilitate calculating contig coverage, gene coverage, and external variant calling. The assembled contigs shorter than a minimal length, or without mapped reads, are filtered out to yield high-quality contigs.

### Genome binning

The prediction of metagenome-assembled genomes (MAGs) allows organism-specific analyses of metagenome datasets. In ATLAS, three binning methods are implemented (Fig. 1): concoct (26), metabat2 (27) and maxbin2 (28). These methods use tetra-nucleotide frequencies, differential abundance, and/or the presence of marker genes as criteria. ATLAS supports assembly and binning for each sample individually, which produces more continuous genomes than co-assembly (29). Definition of which samples are likely to contain the same bacterial species, via a group attribute in the Snakemake configuration file, supports binning based on co-abundance patterns across samples. Reads from all of the samples defined in a group are then aligned to the individual sample assemblies, to obtain the co-abundance patterns needed for efficient binning. The bins produced by the different binning tools can be combined using the dereplicate, aggregate and score tool (DAS Tool, (30)), to yield high-quality MAGs for each sample. Finally, the completeness and contamination of each MAG are assessed using CheckM (31).

Because the same genome may be identified in multiple samples, dRep (29) can be used to obtain a non-redundant set of MAGs for the combined dataset by clustering genomes to a defined average nucleotide identity (ANI, default 0.95) and returning the representative with the highest dRep score in each cluster. dRep first dereplicates using Mash (32), followed by MUMmer (33), thereby benefitting from their combined speed (Mash) and accuracy (MUMmer). The abundance of each genome can then be quantified across samples by mapping the reads to the non-redundant MAGs and determining the median coverage across each the genome.

### Taxonomic and Functional annotation

For annotation, ATLAS supports the prediction of open reading frames (ORFs) using Prodigal (34). The translated gene products are then clustered using linclust (35) to generate non-redundant gene and protein catalogs, which are mapped to the eggNOG catalogue (36) using DIAMOND (37). Robust taxonomic annotation is performed using BAT (38) to map the gene products to a set of non-redundant proteins in GeneBank (39), and infer the taxonomy using a Last Common Ancestor approach based on the high-scoring hits for each gene.

### Output

For each sample, ATLAS produces flat files of the quality-controlled reads, assembled contigs, alignments (bam files), ORF and protein sequences, together with an HTML report containing summary statistics from the quality control, assembly, and genomic binning stages. The genome binning output includes the summary information (quality, abundance) for each genome together with the inferred taxonomy, genome sequence and gene annotations for each of the non-redundant, high-quality MAGs. From the annotation stage, two fasta files are produced containing the nucleotide and amino acid sequences for each gene in the non-redundant gene catalog. The annotations and raw counts for each gene in each sample are provided in a tab-delimited file. Examples of ATLAS output are provided on GitHub (https://github.com/metagenome-atlas) and shown in Fig 2.

**Figure 2.**
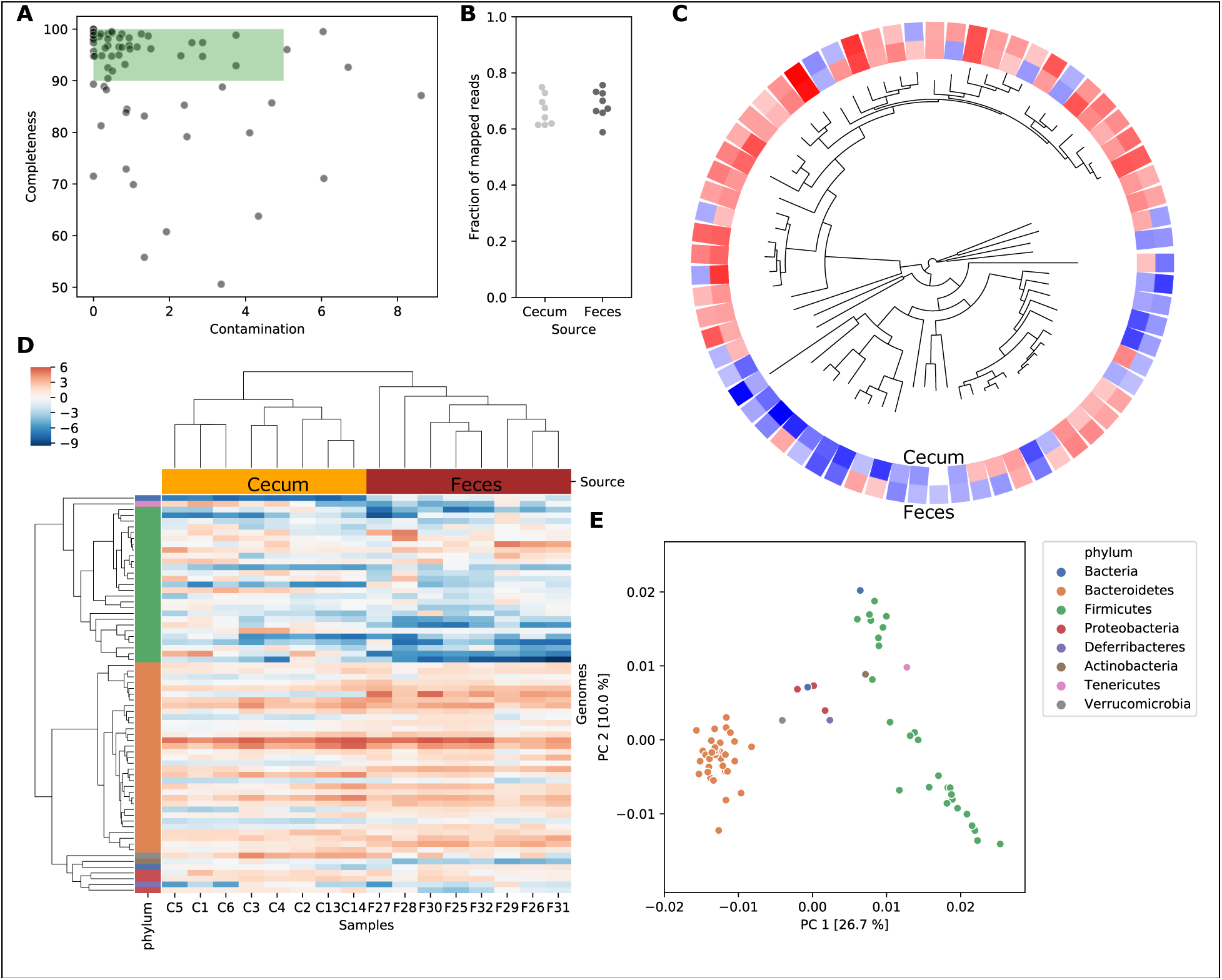
Example output from the ATLAS workflow. Fecal microbiome data (PRJNA480387; (40)) processed by ATLAS show: A) the completeness and contamination of dereplicated MAGs, with high-quality genomes highlighted; B) the fraction of reads mapped to genomes; C) a phylogenetic tree of MAGs with average abundance in feces and cecum on a centered log_2_ scale; D) a heatmap of abundance on a centered log_2_ scale in which MAGs were clustered by phylogenetic distance and samples by Euclidian distance; E) a principle components analysis of the MAGs based on functional annotation.

## Conclusions

ATLAS is easy to install and provides documented and modular workflows for the analysis of metagenome and metatranscriptome data. The internal codes utilized by the workflow are highly configurable using either a configuration file or via the command line. ATLAS provides a robust bioinformatics framework for high-throughput sequence data, where raw FASTQ files can be fully processed into annotated tabular files for downstream analysis and visualization. ATLAS fills a major analysis gap, namely the integration of tools for quality control, assembly, binning and annotation, in a manner that supports robust and reproducible analyses. ATLAS provides these analysis tools in a command-line interface amenable to high-performance computing clusters.

The source code for ATLAS is distributed under a BSD-3 license and is freely available at https://github.com/metagenome-atlas/atlas, with example data provided for testing. Software documentation is available at https://metagenome-atlas.rtfd.io.

## Availability and requirements

Project name: ATLAS

Project home page: https://github.com/metagenome-atlas/atlas

Archived version: https://doi.org/10.7287/peerj.preprints.2843v1

Operating system(s): Linux

Programming language: Snakemake/Python

Other requirements: Miniconda

License: BSD-3

Any restrictions to use by non-academics: None

## Authors’ contributions

JB and SK developed the software and documentation; EZ, MT and LAM supervised the project; and JB, SK and LAM wrote the manuscript. All authors read and approved the final manuscript.

## Acknowledgements

A portion of the ATLAS framework was developed using PNNL Research Computing resources.

